# Induction of Anxiety-Like Phenotypes by Knockout of Cannabinoid Type-1 Receptor in Amygdala of Marmosets

**DOI:** 10.1101/2022.05.24.493231

**Authors:** Di Zheng, Lin Zhu, Rui Li, Chen-Jie Shen, Ruolan Cai, Hao Sun, Xiaohui Wang, Yu Ding, Bin Xu, Guoqiang Jia, Xinjian Li, Lixia Gao, Xiao-Ming Li

## Abstract

The amygdala is an important hub for the regulation of emotions, which is crucial for elucidating cellular and molecular mechanisms of many mental diseases. In the central nervous system, the endocannabinoid system plays a key role in the regulation of emotions and mainly functions through the cannabinoid type-1 receptor (CB1R), which is encoded by the *Cnr1* gene. Although CB1R is highly expressed in the amygdala of non-human primates, little is known about its function. Here, we investigated the function of CB1R by knocking out the CB1R in the amygdala of adult marmosets through regional delivery of AAV-SaCas9-gRNA. We found that CB1R knockout in the amygdala of marmosets induced anxiety-like behaviors, including disrupted night sleep, agitated psychomotor activity in new environments, and reduced social desire, but had no effect on hedonic state and fear response. Moreover, CB1R-knockout marmosets exhibited up-regulated plasma cortisol levels, suggesting increased stress. These results showed that knockout of CB1R in the amygdala induced anxiety-like phenotypes in marmosets and shed new light on the mechanisms underlying the regulation of anxiety by CB1R in the amygdala of non-human primates.

## Introduction

The amygdala is critical for the regulation of emotion (Dunsmoor and Paz, 2015; Gothard, 2020; Janak and Tye, 2015; Morrison and Salzman, 2010) and is implicated in various mental disorders such as depression (Hamilton and Gotlib, 2008), social disturbance (Jayakar et al., 2020), and anxiety (Hyde et al., 2011). As such, it is important that we fully understand its functions.

In the central nervous system, the endocannabinoid system (eCB) functions to guard against negative emotions (Lutz et al., 2015) through the cannabinoid type-1 receptor (CB1R), which is encoded by the *Cnr1* gene and modulates synaptic transmission by suppressing the release of neurotransmitters (Castillo et al., 2012; Ohno-Shosaku and Kano, 2014). CB1R dysfunction is correlated with psychiatric disorders (Choi et al., 2012; Hungund et al., 2004), and manipulation of CB1R in rodents shows bidirectional regulation of anxiety-like behaviors (Haring et al., 2011; Moreira et al., 2009; Rey et al., 2012). CB1R is highly expressed in the amygdala, and associations between CB1R expression in the amygdala and depression-like behavior have been reported in mice (Shen et al., 2019). However, the function of amygdala CB1R in emotional regulation is far less known in non-human primates (NHPs).

As an NHP model organism, marmosets (*Callithrix jacchus*) have garnered considerable interest in studies on emotion and cognition due to their highly vocal and emotionally rich social behaviors (Okano, 2021; Okano et al., 2016).

Marmosets are also invaluable in biomedical research due to their genetic and physiological similarities to humans and relative ease of care in captivity (Marx, 2016). Recent advances in adeno-associated virus (AAV)-mediated delivery of CRISPR/Cas9 in adult macaques (Li et al., 2021; Wu et al., 2021) highlight the potential application of genetically modified adult marmosets in neuroscience. Therefore, in the current study, we explored the function of CB1R in the amygdala of marmosets via *in vivo* gene editing. Through regional delivery of AAV-SaCas9-gRNA, marmosets with CB1R knockout in the amygdala displayed anxiety-like phenotypes. These results revealed the emotion-specific function of the amygdala CB1R in NHPs.

## Results

### Construction and validation of AAV-mediated CRISPR/Cas9 virus

Based on SaCas9 (Hanna and Doench, 2020), an AAV-mediated CRISPR/Cas9 virus was used to knockout the *Cnr1* gene in the amygdala of adult marmosets *in vivo*. To specifically target the *Cnr1* gene, three guide RNAs (gRNAs) were synthesized and inserted into the AAV-hSyn-SaCas9-U6-gRNA vector (Fig. 1A). Considering the deep location and individual variation of the amygdala in NHPs, MRI was applied to localize and target the exact region of the amygdala in marmosets before virus injection (Fig. 1B). To validate knockout efficiency of the virus, one marmoset (Ctrl 1) receiving control virus injection and one marmoset (S) receiving knockout (KO) virus injection were sacrificed via transcardial perfusion, and brain slices were collected for fluorescence staining to evaluate knockout efficiency. Using RNAscope staining, we found much fewer cells expressing CB1R mRNA in the amygdala of KO marmoset than control marmoset (Fig. 1C-E). The numbers of GFAP+ cells and Iba1+ cells (markers of astrocytes and microglial cells, respectively) were the same in control and KO marmosets (Figure 1-figure supplement 1). These results indicate that the AAV-mediated CRISPR/Cas9 *in vivo* gene editing successfully knocked out the *Cnr1* gene in the amygdala of adult marmosets without difference of neuroinflammation effects between two groups.

**Fig. 1:**
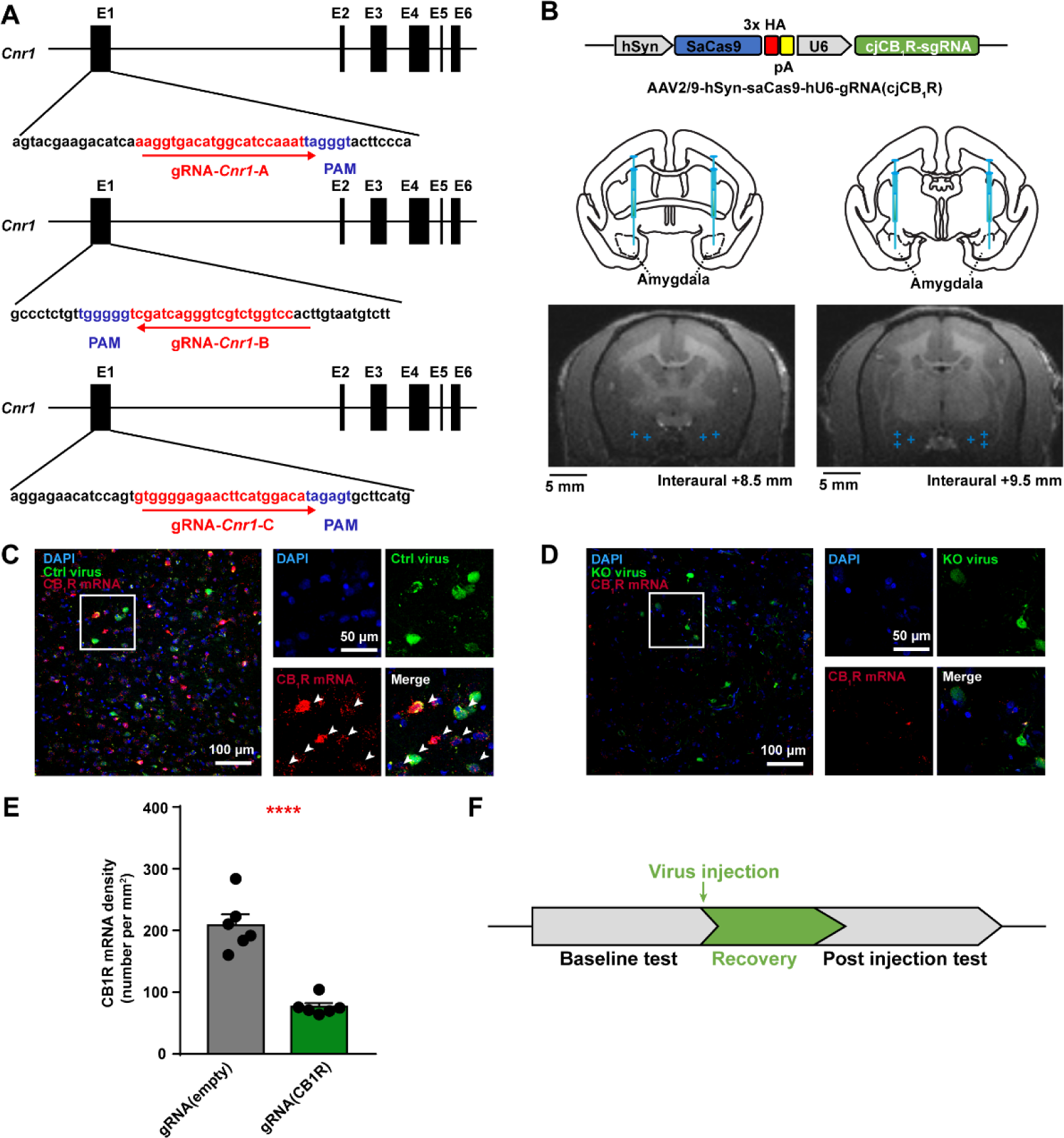
Construction, injection, and validation of AAV-mediated SaCas9 viruses in amygdala of adult marmosets. **A**, Targeting sites of gRNA for marmoset *Cnr1* gene (red). PAM sequences are indicated in blue. **B**, Top, schematic of KO virus; Middle, atlas of marmoset brain with injection sites in amygdala; Bottom, examples of MRI images with amygdala injection sites (indicated with blue cross). **C**, **D**, Left, RNAscope staining of CB1R mRNA (indicated by white arrows) in the amygdala of control (**C**) and KO (**D**) marmoset; Right, magnified view of rectangle on left. Scale bar, 100 μm (left); 50 μm (right). **E**, Density of CB1R mRNA-expressing cells in amygdala. n = 6 slices, two-tailed unpaired *t*-test, t(10) = 7.209, *P* < 0.0001. **F**, Workflow of experiments. **** *P* < 0.0001; data are mean ± SEM. Figure 1-figure supplement 1 was related. Source data were provided in figure 1-source data 1 and 2.

Based on virus injection differences, six marmosets were divided into control and KO groups for behavioral tests, with two males and one female in each group (Table 1). To better investigate the effect of CB1R knockout in the amygdala, we compared behaviors individually before and two months after virus injection (Fig. 1F). Comparisons of changes in behavior between the two groups were also conducted.

**Table 1.** Detailed information on marmosets used in study.

| Group | ID | Age | Sex | Virus injected |
| --- | --- | --- | --- | --- |
| Control | Ctrl1 | 5 | Male | gRNA(empty) |
| Control | Ctrl2 | 3 | Male | gRNA(empty) |
| Control | Ctrl3 | 3 | Female | None |
| KO | KO1 | 4 | Male | gRNA(cjCB<sub>1</sub>R) |
| KO | KO2 | 3 | Male | gRNA(cjCB<sub>1</sub>R) |
| KO | KO3 | 3 | Female | gRNA(cjCB<sub>1</sub>R) |
| KO | S | 3 | Male | gRNA(cjCB<sub>1</sub>R) |

### Disrupted night sleep in CB1R knockout marmosets

As many emotional disorders co-occur with sleep disturbance (Qiu et al., 2019), we monitored the sleep of marmosets using an actigraphy device (Paquet et al., 2007; Ross et al., 2019; Zhou et al., 2019). Results showed that sleep latency to light-off in control marmosets remained the same (Fig. 2A, B, and D) but was prolonged in KO marmosets with an increasing trend (Fig. 2F, G, and I) after virus injection. In addition, night sleep duration in control marmosets did not change (Fig. 2C and E) but was shortened in KO marmosets with a decreasing trend (Fig. 2H and J) after virus injection. By comparing changes in sleep latency and sleep duration between groups, we also found prolonged sleep latency (Fig. 2K) and reduced sleep duration (Fig. 2L) in the KO group animals. We also examined whether decreased night sleep was due to increased daytime sleep but found that daytime nap time was not influenced by virus injection in either the control or KO groups (Figure 2—figure supplement 1). We also explored sleep efficiency (Allison et al., 2016) and fragment index (Zhou et al., 2019), which both contribute to sleep quality, and found that neither were affected in control marmosets (Figure 2—figure supplement 2A and B). However, sleep efficiency showed a decreasing trend (Figure 2—figure supplement 2C and E) and the fragment index showed an increasing trend (Figure 2—figure supplement 2D and F) in KO marmosets, suggesting decreased sleep quality.

**Fig. 2:**
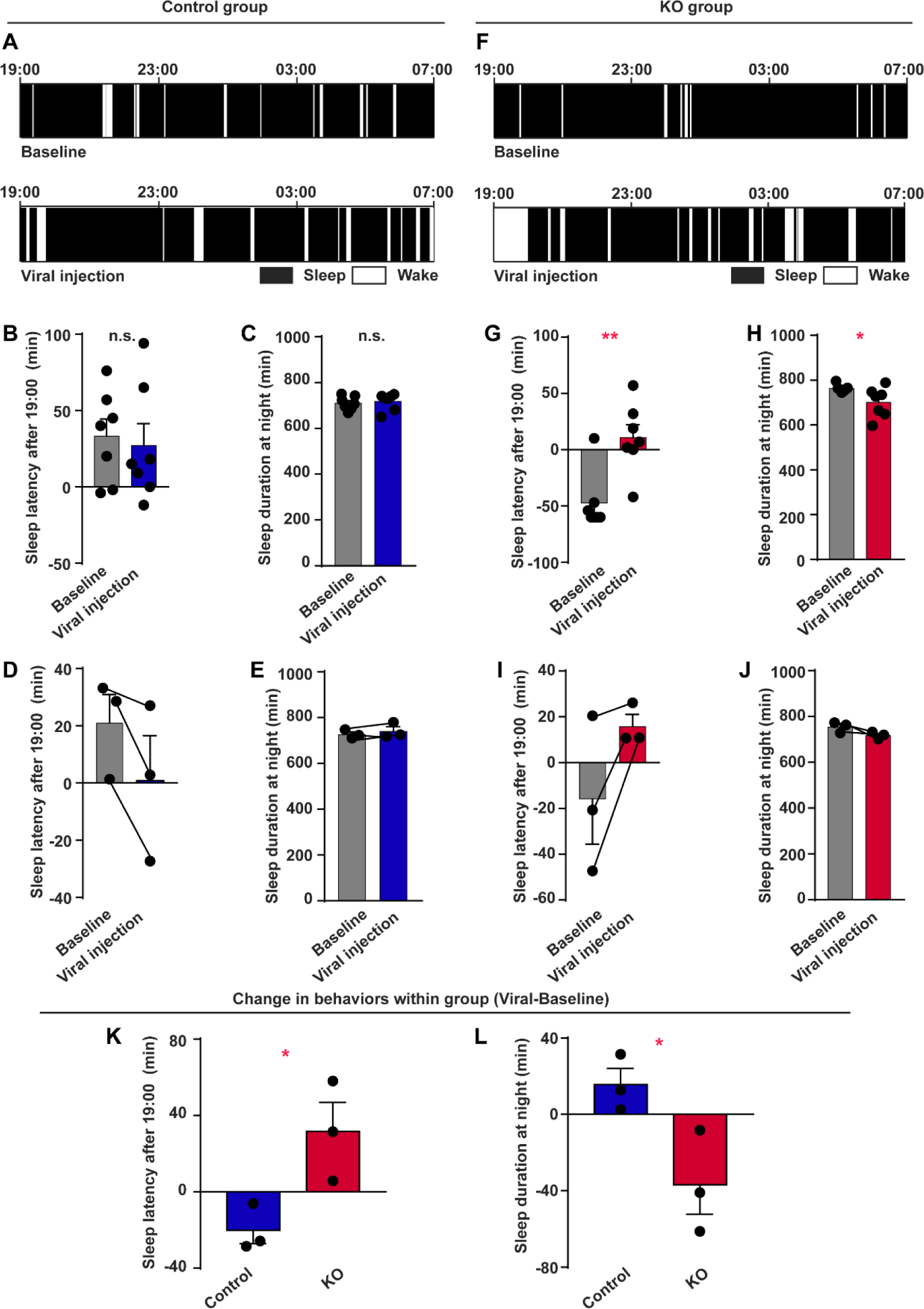
Disrupted night sleep in CB1R knockout marmosets. **A**, **F**, Representation of sleep patterns at night (19 PM to 7 AM) before virus injection (top) and after virus injection (bottom) in a control (**A**) and a KO (**F**) marmoset. Sleep phase is indicated by black box, awake phase is indicated by white box. **B**, **G**, Sleep latency in a control (**B**) and a KO (**G**) marmoset. Mann-Whitney *U* test, U = 21, *P* = 0.7104 (**B**); U = 4, *P* = 0.0064 (**G**). **C**, **H**, Night sleep duration in a control (**C**) and a KO (**H**) marmoset. Mann-Whitney *U* test, U = 22, *P* = 0.8048 (**C**); U = 7, *P* = 0.0239 (**H**). **D**, **I**, Comparison of 7-day average sleep latency before and after virus injection for individuals in control (**D**) and KO (**I**) groups. n = 3, two-tailed paired *t*-test, t(2) = 2.858, *P* = 0.1037 (**D**); t(2) = 2.104, *P* = 0.1700 (**I**). **E**, **J**, Comparison of 7-day average night sleep duration for individuals in control (**E**) and KO (**J**) groups. n = 3, two-tailed paired *t*-test, t(2) = 1.529, *P* = 0.2659 (**E**); t(2) = 2.387, *P* = 0.1396 (**J**). **K**, Comparison of changes in sleep latency between control and KO groups. n = 3, two-tailed unpaired *t*-test, t(4) = 3.116, *P* = 0.0357. **L**, Comparison of changes in night sleep duration between control and KO groups. n = 3, two-tailed unpaired *t*-test, t(4) = 2.832, *P* = 0.0472. * *P* < 0.05; data are mean ± SEM. Figure 2-figure supplement 1 and 2 were related. Source data were provided in figure 2-source data 1 to 3.

Taken together, our results suggest that knockout of CB1R in the amygdala disrupts night sleep quantity and quality in marmosets, implicating emotional dysregulation.

### Agitated psychomotor activity in new environment in CB1R knockout marmosets

New environments are stressful and may induce behavioral and emotional abnormalities in animals and humans (Penninx et al., 2021; Spasojevic et al., 2016). We transferred single marmosets to a new environment (cage) (Qiu et al., 2019) to test the effects on behavior. In contrast to the control group, the KO marmoset trajectories were much denser after virus injection, indicating agitation in the new environment (Fig. 3A and E). In addition, while moving distance, average velocity, and stationary time remained the same in the control group (Fig. 3B–D), KO marmosets showed increased moving distance and average velocity (Fig. 3F and G) as well as decreased stationary time (Fig. 3H) after virus injection. Comparisons between groups further confirmed higher agitation in the KO marmosets, with increased moving distance (Fig. 3I) and average velocity (Fig. 3J) and decreased stationary time (Fig. 3K). Marmoset activity in their home cages was also measured by actigraphy (Figure 3—figure supplement 1A). However, no significant changes were found in either group (Figure 3—figure supplement 1B–F), indicating that motor ability was unaffected by CB1R knockout in the amygdala.

**Fig. 3:**
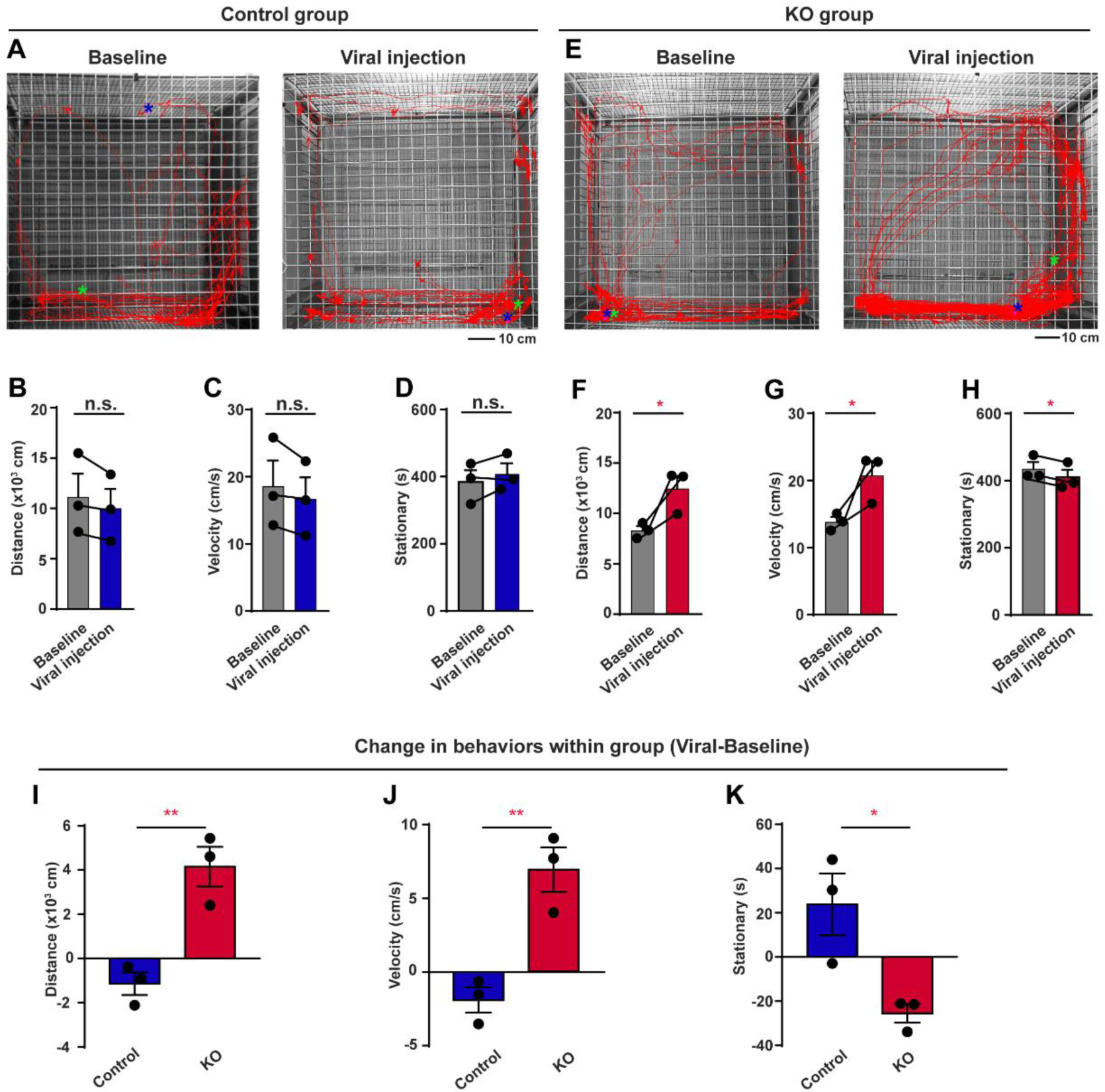
Agitated psychomotor activity in new environment for CB1R knockout marmosets. **A**, **E**, Representation of moving trajectories in new environment before virus injection (left) and after virus injection (right) in a control (**A**) and a KO (**E**) marmoset. Start and end positions are indicated by green and blue asterisks, respectively. Scale bar, 10 cm. **B**, **F**, Seven-day average travel distance in new environment before and after virus injection in control (**B**) and KO (**F**) groups. N = 3, two-tailed paired *t*-test, t(2) = 2.227, *P* = 0.1558 (**B**); t(2) = 4.577, *P* = 0.0446 (**F**). **C**, **G**, Seven-day average velocity in new environment before and after virus injection in control (**C**) and KO (**G**) groups. n = 3, two-tailed paired *t*-test, t(2) = 2.227, *P* = 0.1558 (**C**); t(2) = 4.577, *P* = 0.0446 (**G**). **D**, **H**, Seven-day average stationary time in new environment before and after virus injection in control (**D**) and KO (**H**) groups. n = 3, two-tailed paired *t*-test, t(2) = 1.703, *P* = 0.2307 (**D**); t(2) = 6.078, *P* = 0.0260 (**H**). **I-K**, Comparison of changes in travel distance (**I**), average velocity (**J**), and stationary time (**K**) in new environment between control and KO groups. n = 3 marmosets in each group, two-tailed unpaired *t*-test, t(4) = 5.080, *P* = 0.0071 (**I**); t(4) = 5.080, *P* = 0.0071 (**J**); t(4) = 3.379, *P* = 0.0278 (**K**). * *P* < 0.05; ** *P* < 0.01; n.s., not significant; data are mean ± SEM. Figure 3-figure supplement 1 was related. Source data were provided in figure 3-source data 1 and 2.

The above results suggest that CB1R knockout in the amygdala of marmosets specifically induces agitated psychomotor activity in stressful new environments but not in familiar surroundings.

### Decreased social desire for vocal communication in CB1R knockout marmosets

Marmosets are highly vocal and social animals (Miller et al., 2016) and use phee calls to communicate with each other (Bezerra and Souto, 2008). We hypothesized that CB1R knockout in the amygdala may induce abnormal social behaviors in marmosets, reflected by a decrease in vocal production. We recorded phee calls of separated marmosets to test social behavior. Interestingly, while phee calls did not change in control marmosets (Fig. 4A and B), the KO marmosets tended to produce fewer phee calls after virus injection, indicating less vocal social behavior (Fig. 4C and D). Intergroup comparison showed that knockout of CB1R resulted in a decrease in social phee calls (Fig. 4E), implying a reduction in vocal social desire.

**Fig. 4:**
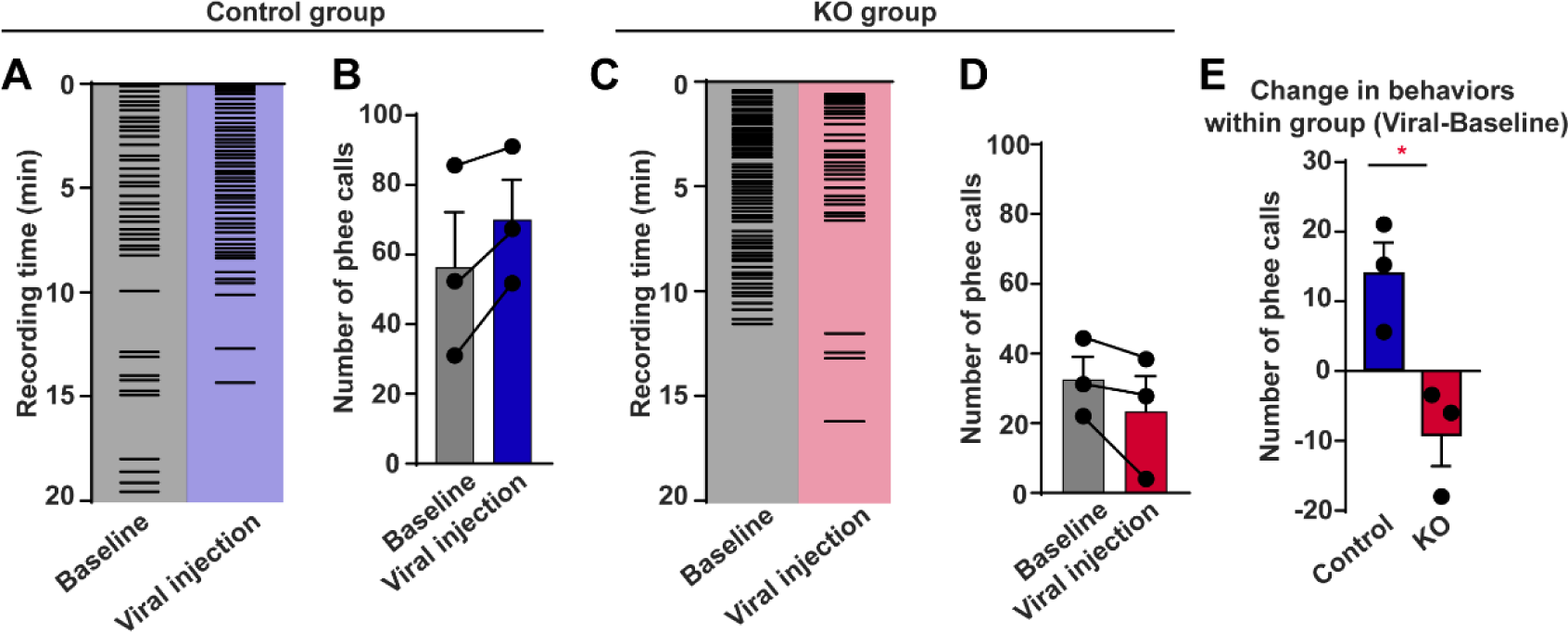
Decreased social desire for vocal communication in CB1R knockout marmosets. **A**, **C**, Example raster plots of phee calls during vocal recordings before virus injection and after virus injection in a control (**A**) and a KO (**C**) marmoset. **B**, **D**, Five-day average number of phee calls before and after virus injection in control (**B**) and KO (**D**) groups. n = 3, two-tailed paired *t*-test, t(2) = 3.058, *P* = 0.0923 (**B**); t(2) = 0.1793, *P* = 2.031 (**D**). **E**, Comparison of changes in phee calls between control and KO groups. n = 3 marmosets in each group, two-tailed unpaired *t*-test, t(4) = 3.598, *P* = 0.0228. * *P* < 0.05; data are mean ± SEM. Figure 4-figure supplement 1 was related. Source data were provided in figure 4-source data 1 and 2.

We also assessed social behavior using the three-chamber social test, as used in rodents (Rein et al., 2020). After habituation, marmosets were tested by the introduction of a stranger followed by the introduction of a cage-mate (Figure 4—figure supplement 1A). However, no significant changes in social behavior were observed in either the control or KO groups (Figure 4—figure supplement 1B–D). A potential explanation may be that rodent behavioral tests are unsuitable for testing sociability in NHPs, given that different species use different modalities to recognize and communicate with conspecifics (Miller et al., 2016).

Taken together, our results suggest that knockout of CB1R in the amygdala decreases social desire in marmosets, implicating anxiety-like social avoidance.

### Unaltered hedonic and fear behaviors in CB1R knockout marmosets

We previously showed that disruption of circuit-specific CB1R in the amygdala induces depression-like anhedonia in mice (Shen et al., 2019). In the current study, we tested the hedonic state in marmosets via an adapted sucrose preference test (Alexander et al., 2019) (Fig. 5A). In the test, marmosets consumed sucrose and water gradually over 2 h (Figure 5—figure supplement 1) and sucrose preference was examined in the first 30 and 120 min, respectively. Results showed that sucrose preference was not changed after virus injection in either the control or KO groups (Fig. 5B–D), suggesting no depression-like behavior.

**Fig. 5:**
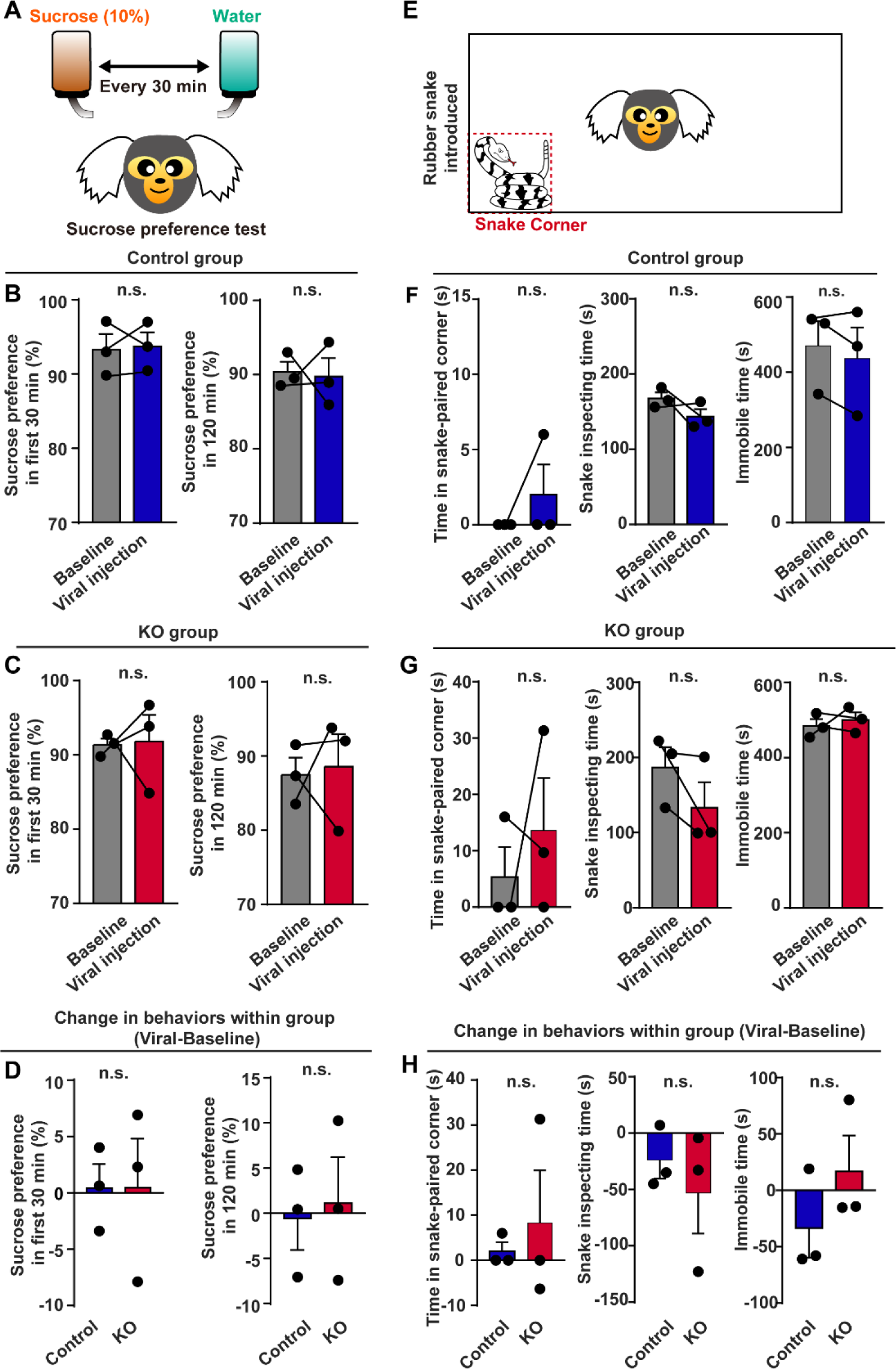
Unaltered hedonic state and fear response in CB1R knockout marmosets. **A**, Paradigm of sucrose preference test. **B**, **C**, Individual sucrose preference in first 30 min (left) and 120 min (right) before and after virus injection in control (**B**) and KO (**C**) groups. n = 3, two-tailed paired *t*-test, t(2) = 0.2001, *P* = 0.8599 (**B**, left); t(2) = 0.1724, *P* = 0.8790 (**B**, right); t(2) = 0.1042, *P* = 0.9265 (**C**, left); t(2) = 0.2166, *P* = 0.8486 (**C**, right). **D**, Comparison of changes in sucrose preference in first 30 min (left) and 120 min (right) between control and KO groups. n = 3, two-tailed unpaired *t*-test, t(4) = 0.0057, *P* = 0.9957 (left); t(4) = 0.2760, *P* = 0.7962 (left)**. E**, Paradigm of snake intruder test. **F**, **G**, Individual time spent in snake corner (left), time spent inspecting snake (middle), and immobile time (right) before and after virus injection in control (**F**) and KO (**G**) groups. n = 3, two-tailed paired *t*-test, t(2) = 1.000, *P* = 0.4226 (**F**, left); t(2) = 1.527, *P* = 0.2662 (**F**, middle); t(2) = 1.273, *P* = 0.3309 (**F**, right); t(2) = 0.7155, *P* = 0.5486 (**G**, left); t(2) = 1.492, *P* = 0.2742 (**G**, middle); t(2) = 0.5324, *P* = 0.6476 (**G**, right). **H**, Comparison of changes in time spent in snake corner (left), time spent inspecting snake (middle) and immobile time (right) between control and KO groups. n = 3, two-tailed unpaired *t*-test, t(4) = 0.5368, *P* = 0.6198 (left); t(4) = 0.7391, *P* = 0.5009 (middle); t(4) = 1.222, *P* = 0.2889 (left). n.s., not significant; data are mean ± SEM. Figure 5-figure supplement 1 was related. Source data were provided in figure 5-source data 1.

Previous studies have indicated that the amygdala is a pivotal region for fear response (Gothard, 2020; Janak and Tye, 2015). Thus, we measured fear response in marmosets using the snake intruder test based on previous protocols (Melamed et al., 2017) (Fig. 5E). Unexpectedly, knockout of CB1R in the amygdala did not change fear response behavior in marmosets (Fig. 5F– H).

The above results suggest that knockout of CB1R in the amygdala did not change hedonic state or innate fear response in marmosets.

### Increased plasma cortisol level in CB1R knockout marmosets

Our results showed that CB1R knockout produced anxiety-like behaviors in marmosets, which may induce *in vivo* biochemical abnormalities, such as increased plasma cortisol (Chida and Steptoe, 2009). Thus, we tested whether behavioral disruptions were accompanied by abnormal cortisol levels. Morning blood samples were obtained from marmosets, followed by plasma collection and ELISA analysis (Fig. 6A). Interestingly, plasma cortisol levels showed an increasing tendency in the KO group but not in the control group after viral injection (Fig. 6B and C). Furthermore, intergroup comparison revealed that plasma cortisol levels were higher in the KO marmosets than in the controls (Fig. 6D).

**Fig. 6:**
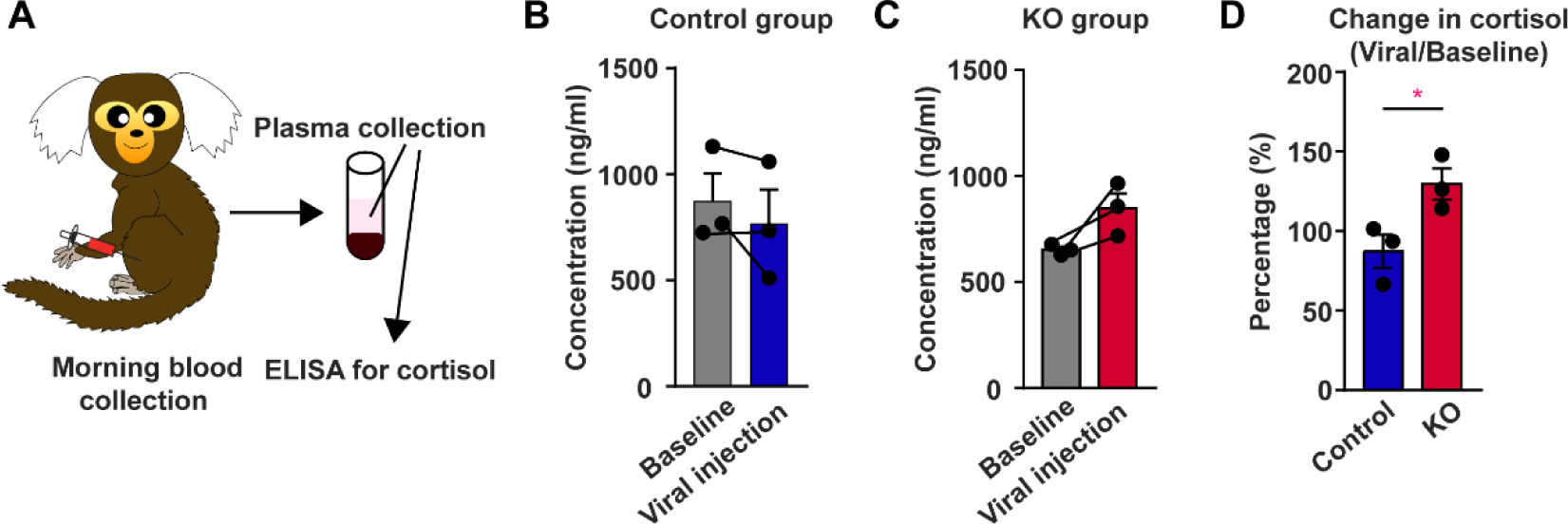
Increased plasma cortisol level in CB1R knockout marmosets. **A**, Paradigm of plasma cortisol level evaluation. **B**, **C**, Individual plasma cortisol level before and after virus injection in control (**B**) and KO (**C**) groups. n = 3, two-tailed paired *t*-test, t(2) = 1.346, *P* = 0.3106 (**B**); t(2) = 2.990, *P* = 0.0960 (**C**). **D**, Comparison of changes in plasma cortisol level between control and KO groups. n = 3, two-tailed unpaired *t*-test, t(4) = 2.933, *P* = 0.0427. * *P* < 0.05; data are mean ± SEM. Source data were provided in figure 6-source data 1.

These results indicate that CB1R knockout in the amygdala induces anxiety-like behaviors in marmosets accompanied by increasing plasma cortisol levels.

## Discussion

Using *in vivo* gene editing, immunohistochemical analysis, behavioral testing, and biochemical measurement, we studied the function of CB1R in the amygdala of adult marmosets. Results showed that AAV-mediated delivery of the CRISPR/Cas9 system successfully knocked out CB1R expression in the amygdala of adult marmosets and induced anxiety-like phenotypes (Park and Kim, 2020; Penninx et al., 2021), including disrupted night sleep, agitated psychomotor activity in new environments, decreased desire for vocal communication, and increased plasma cortisol levels. Applying gene editing to study the region-specific functions of CB1R in adult marmosets should help improve our understanding of the eCB system in the amygdala of NHPs and advance the clinical application of eCB for the diagnosis and treatment of psychiatric disorders, such as anxiety.

Although rodents are useful models in neuroscience, they differ from humans in multiple aspects, which may hinder our understanding of the mechanisms underlying human emotion, cognition, and mental disorders (Camus et al., 2015; Feng et al., 2020; Izpisua Belmonte et al., 2015). As such, studying the brains of NHPs is critical for translational research due to their homogeneity to humans (Jennings et al., 2016). Electrolytic lesions and pharmacological manipulations via cannulas are commonly used to explore the general function of the amygdala in NHPs (Braesicke et al., 2005; Dal Monte et al., 2015; Wellman et al., 2016), but cause non-negligible side effects and lack molecular marker specificity. At present, gene editing techniques widely used in rodents are not well established in NHPs, which greatly impedes our understanding of the brain mechanisms that underpin mental disorders in humans. Here, for the first time, we established an *in vivo* gene editing method to study the function of CB1R in the amygdala of marmosets. We successfully demonstrated the feasibility of virus-mediated gene editing in adult NHPs, thus providing a novel strategy to develop transgenic NHP models in a rapid and cost-effective manner (Marx, 2016).

Following CB1R knockout in the amygdala, marmosets showed various anxiety-related behaviors, consistent with previous reports indicating that the amygdala and CB1R are involved in the onset and development of anxiety (Janak and Tye, 2015; Lutz et al., 2015; Ramikie and Patel, 2012). Of note, we detected sleep disturbance and agitation in the marmosets with anxiety-like behaviors, which are rarely detected in anxiety model rodents but are key pathological manifestations of anxiety in humans (Park and Kim, 2020). We also detected increased plasma cortisol levels in the KO marmosets, consistent with previous study (Akirav, 2013) and a representative biomarker of anxiety in humans (Chida and Steptoe, 2009), but which is less noted in rodent experiments (Calhoon and Tye, 2015; Tovote et al., 2015). Notwithstanding, the causal relationship between the change in cortisol level and behavior remains unclear and deserves further investigation. However, our results suggest that NHP models are crucial for studying human-relevant diseases and exhibit considerable potential in identifying specific molecular biomarkers for the diagnosis and treatment of psychiatric disorders.

As CB1R exerts its function through synaptic modulation, and not all neurons in the amygdala express CB1R (Shen et al., 2019), a potential explanation for our findings may be that neurons that regulate anxiety state may express higher levels of CB1R. Thus, knockout of CB1R may disrupt the anxiogenic-anxiolytic balance, thereby potentiating anxiety transmission by enhancing amygdala-related anxiogenic circuit activity. However, further circuit-based studies are required to confirm this hypothesis.

In contrast to previous studies in rodents (Roche et al., 2007; Shen et al., 2019), we found that CB1R knockout in marmosets did not result in depression-like behaviors or altered fear response, which may be explained by species differences. In addition, the previous rodent-based CB1R knockout studies were circuit specific, cell-type specific, or amygdala-subregion specific, whereas we targeted the entire amygdala region regardless of circuit or cell type. Furthermore, as CB1R is mainly distributed in the presynaptic membrane (Lutz et al., 2015), the intra-amygdala infusion of CB1R agonist and antagonist in previous research may have affected CB1R in the amygdala-projecting terminals rather than the amygdala itself, which may complicate the effects. As such, the underlying mechanism still awaits further investigation.

There are some limitations in the current study. As mentioned above, we treated the amygdala as a whole and different types of cells and subregions in the amygdala may compensate in function, leading to less dramatic changes in marmoset behavior. Thus, future studies on the different subregions and types of neurons in the NHP amygdala are needed. The use of Cre-expressing marmosets may also be insightful (Okano, 2021). The small sample size in our study due to animal number limitations during the COVID-19 pandemic may explain why some behavioral trends did not reach significance. However, the significant behavioral changes in the KO group were considered prominent and robust.

In conclusion, we established a novel gene editing method in adult marmosets and highlighted the function of CB1R in the marmoset amygdala in the regulation of anxiety-like phenotypes, thus paving the way for further use of NHPs in basic research.

## Materials and Methods

### Animal ethics

Male and female marmosets (350–450 g, 2–4 years old) were purchased from Johnbio (Jiangsu, China) and kept in the Non-Human Primate Center of Zhejiang University (ZJU) in pairs. All experiment protocols were conducted under the guidelines for the care and use of laboratory animals of ZJU and were approved by the Animal Advisory Committee at ZJU following the National Institutes of Health (NIH) guidelines. Animal details are provided in Table 1.

### Viruses

All viruses were constructed and purchased from Taitool Bioscience (Shanghai, China). AAV2/9-hSyn-saCas9-hU6-gRNA(cjCB1R) (1.21 × 10^13^ viral genomes (vg)/ml) was used to knockout the *Cnr1* gene, AAV2/9-hSyn-saCas9-pA-hU6-gRNA(empty) (1.58 × 10^13^ vg/mL) was used as the control virus, and AAV2/9-hSyn-EGFP-WPRE-pA (1.42 × 10^13^ vg/mL) was co-injected with the above viruses (1:9) to display injection location.

### Magnetic resonance imaging (MRI)-guided amygdala location

MRI data were acquired on a 7-T research scanner (Siemens Healthcare, Erlangen, Germany) with a single loop coil (RAPID MR International, Columbus, OH, USA) for signal reception and transmission. Structural images were obtained at an isotropic voxel size of 0.3 mm. Briefly, animals were anesthetized with alfaxalone (intramuscular (i.m.), 0.1 mg/kg, Jurox, North Kansas City, USA) and maintained by sustained isoflurane (0.5%–1.2%, Yipin Pharmacology, Hebei, China). Two glass tubes (OD: 1.0 mm, ID: 0.58 mm; Sutter Instrument, USA) filled with gel were placed interaurally as markers. After the animal was fixed in a home-made MRI compatible stereotaxic apparatus, the scan procedure was started. The MRI data were analyzed using MRIcron v1.0 (NITRC, USA) to calculate coordinates of the amygdala to guide virus injection.

### Virus injection

The surgery and virus injection were performed under aseptic conditions. The animals were initially anesthetized with alfaxalone (i.m., 0.1 mg/kg, Jurox, North Kansas City, USA) and maintained by isoflurane (0.5%–1.2%, Yipin Pharmacology, Hebei, China). The head was fixed in a stereotaxic frame (RWD, Shenzhen, China) and a craniotomy was carried out using a dental drill according to the coordinates. A cocktail of viruses was infused through a Hamilton syringe (Hamilton, USA) placed in a syringe pump (KD Scientific, USA) at a speed of 80 nL/min for a total of 1 μL at each injection site. After infusion, the syringe was left in place for 5 min. The same operation was repeated until all coordinates were injected. After injection, the wound was carefully cleaned and sutured. The animals were given special care for two weeks with additional nutritional supplements and physiological examination. Post-injection measurements were performed two months after injection.

### Histological analysis and imaging

Animals were first euthanized by administering an overdose of sodium pentobarbitone (100 mg/kg) and then transcardially perfused with phosphate-buffered saline (PBS), followed by 4% paraformaldehyde. Brains were removed and post-fixed in 4% paraformaldehyde overnight at 4 °C, and then immersed in 30% sucrose in PBS for 72 h. The brains were embedded in Optimal Cutting Temperature compound (Sakura Finetek, USA) and 20-μm cryosections were cut using a cryostat (Leica Microsystems, Germany), then washed with PBS (5 min/time).

RNAscope *in-situ* hybridization was performed using a RNAscope Multiplex Fluorescent Reagent Kit v2 (ACDbio, USA) and RNAscope Probe-cj-*Cnr1* (Cat No. 565041) (ACDbio, USA) according to the manufacturer’s standard protocols.

For immunofluorescence staining, sections first underwent target retrieval pretreatment, and were then blocked with 3% bovine serum albumin in PBST (0.3% Triton X-100 in PBS) for 1 h and incubated with primary antibodies overnight at 4 °C. The sections were then incubated with fluorescent secondary antibodies (1:400, Invitrogen, USA) for 2 h at room temperature. The primary antibodies were anti-GFAP (1:800, BioLegend, SMI-21R, USA) and anti-Iba1 (1:800, Wako, 019-19741, Japan). Anti-GFP was used to better show EGFP expression in the amygdala. Sections were mounted after 4′,6-diamidino-2-phenylindole (DAPI) staining (1:5 000, Sigma-Aldrich, USA). Confocal images were captured under 20× objective using A-1R Confocal Microscope (Nikon, Japan). Cell counts were performed with ImageJ v1.52 (NIH, USA).

### Daily activity and sleep measurement in home cages

Daily activity and sleep patterns were assessed using an ActiWatch Mini (CamNtech, UK) worn on the neck of each animal. Animals were allowed to habituate to the device for 2 days followed by 7-day data collection. Daytime was defined as 07:00 am to 19:00 pm and night-time was defined as 19:00 pm to 07:00 am according to light/dark cycle (lights on at 07:00 am and off at 19:00 pm). Activity data were analyzed in 1-min epochs using Sleep Analysis v7 (CamNtech, UK) off-line. Sleep latency was defined by the timepoint of the animal falling asleep relative to 19:00 pm. Sleep duration was defined as the time difference between light off and the time the animal fell asleep. Sleep efficiency was defined as the percentage of sleep time minus waking time. Sleep fragment index was determined by the percentage of waking time plus the percentage of immobile time. Calculation details are provided in the Sleep Analysis v7 software guidelines.

### New environment activity examination

The marmosets were transferred to a new cage (1 × 1 × 1 m) and their activity was recorded with a camera for 10 min. Trajectories and moving distances were analyzed using a custom Matlab 2017b (MathWorks, USA) program (provided by Prof. Xinjian Li). Average velocity was defined as moving distance divided by recording time. Stationary time was defined as total duration of immobile phases of more than 1 s.

### Three-chamber social test

The three-chamber social test was modified from that used for mice and behavior was recorded by camera. Briefly, marmosets were first habituated to the three-chamber test apparatus for 20 min. A non-cage-mate stranger marmoset (opposite sex) was then introduced to the left or right chamber for 20 min (stranger introduced session), followed by a cage-mate introduction to the chamber on the other side (not the central chamber) for 20 min (cage-mate introduced session). Every day, the positions of the stranger and cage-mate were counterbalanced in limit place preference. Time spent in each chamber in the two sessions was analyzed manually.

### Vocal test

The animals were separated from their home cage-mates and colony and transferred to the testing room. A microphone was placed in front of the transfer cage for vocal recording for 20 min. Phee calls were manually analyzed using Adobe Audition CS6 v5.0 (Adobe, USA).

### Sucrose preference test

Sucrose preference tests were performed in the home cages in the morning. Before testing, cage-mates were transferred to an adjacent cage, after which, each marmoset was provided with two bottles containing 100 mL of 10% sucrose solution and 100 mL of water, respectively. The sucrose preference test lasted for 2 h. The weights of the sucrose and water were measured and the positions of the two bottles were exchanged every 30 min. Sucrose preference index was V_sucrose consumed_ / (V_sucrose consumed_ + V_water consumed_).

### Snake intruder test

Animals were first habituated to the test apparatus for 10 min. A rubber snake was then introduced in the corner and monkey behavior was recorded by video. Snake-paired corner time was determined as the time spent in the same corner as the snake. Snake inspection time was determined as the time spent staring at the snake. Immobile time was determined as the time spent immobile for more than 1 s.

### Blood sample collection and cortisol test

Blood samples (0.5 mL) were collected intravenously in EDTA-coated tubes at 10:00 am for two continuous days. The blood samples were immediately centrifuged at 1 000 ×*g* for 10 min at room temperature. The plasma was collected and stored at −80 °C until use. Plasma cortisol levels were measured using an ELISA toolkit (Enzo, ADI-901-071, USA) according to the standard protocols provided.

### Quantification and statistical analysis

Sample size for statistical comparisons were referred to previous research (Li et al., 2021; Wu et al., 2021). All data were analyzed with GraphPad Prism v6.01 (GraphPad Software, USA). Differences in individuals before and after virus injection were tested using two-tailed paired *t*-test and differences between groups were tested using two-tailed unpaired *t*-test. Repeated two-way analysis of variance (ANOVA) was used to test differences in the three-chamber test. Data are shown as mean ± standard error of the mean (SEM). Differences were considered statistically significant at * *P* < 0.05, ** *P* < 0.01, *** *P* < 0.001, and **** *P* < 0.0001.

All data generated or analyzed during this study are included in the manuscript and supporting file; Source Data files have been provided.

## Acknowledgements

We thank Dr. Zilong Qiu for providing the AAV-SaCas9 vector. We are grateful to Research Assistant Shuangshuang Liu from the Core Facilities of Zhejiang University School of Medicine, as well as Dr. Sanhua Fang and Research Assistant Li Liu from the Core Facilities of Zhejiang University Institute of Neuroscience. This work was supported by Zhejiang Province Natural Science Foundation of China (LD22H090003), Key-Area Research and Development Program of Guangdong Province (2019B030335001 and 2018B030334001), National Natural Science Foundation of China (31871070, 82090031, 32071097, 31871056 and 32170991), Key R&D Program of Zhejiang Province (2020C03009), Fundamental Research Funds for the Central Universities (2021FZZX001-37), Non-Profit Central Research Institute Fund of the Chinese Academy of Medical Sciences (2019PT310023), and CAMS Innovation Fund for Medical Sciences (2019-I2M-5-057).

## Figure Supplements

**Figure 1—figure supplement 1.**
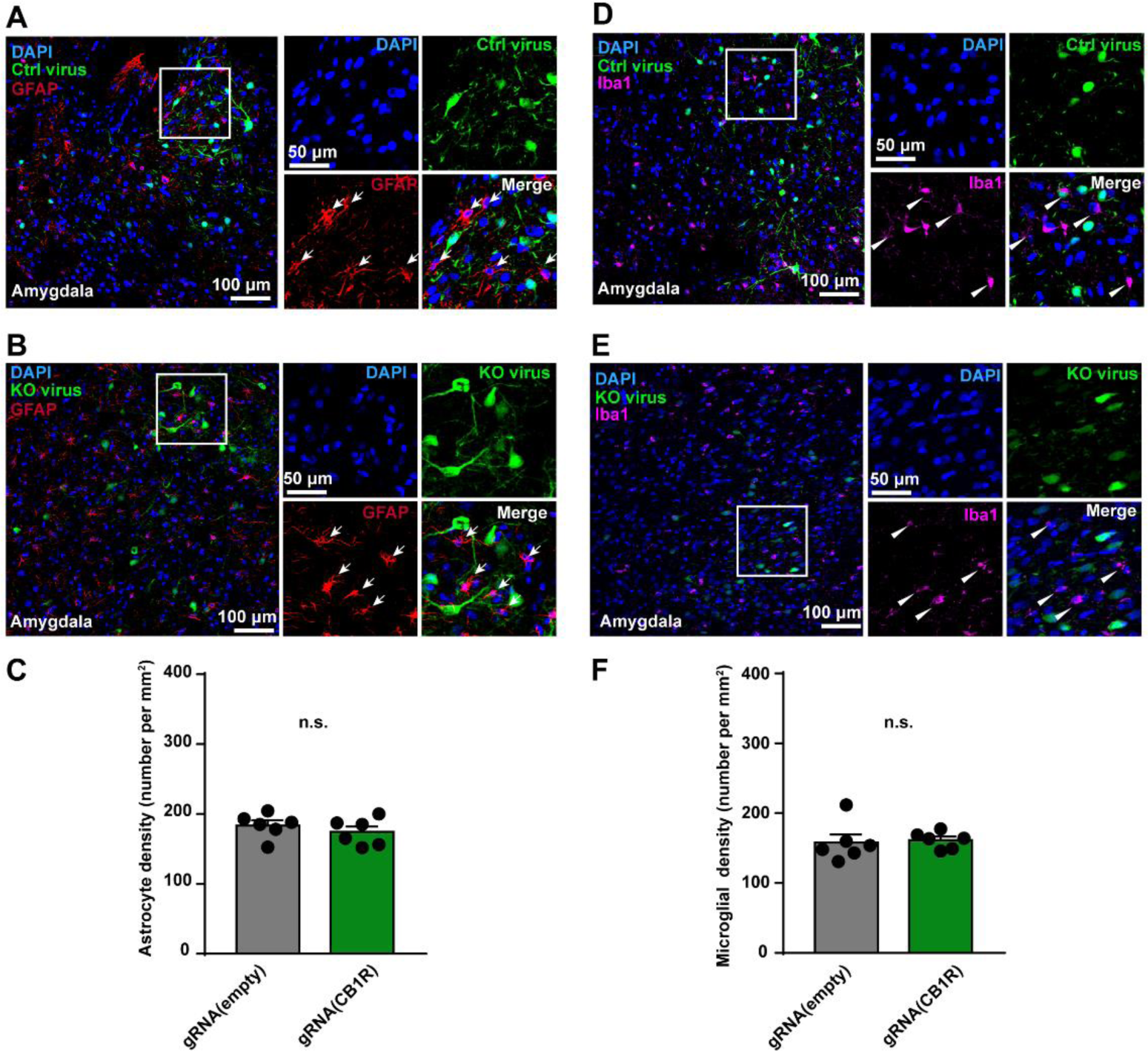
Immunohistochemical analysis of astrocytes and microglial cells in the amygdala of marmosets. **A**, **B**, Left, GFAP-positive astrocytes (red, indicated by arrows) in the amygdala of control (**A**) and KO (**B**) marmosets. Right, magnified view of rectangle on left. Scale bar, 100 μm (left); 50 μm (right). **C**, Density of astrocytes in the amygdala of marmosets. n = 6 slices, two-tailed unpaired *t*-test, t(10) = 0.8672, *P* = 0.4062;. **D**, **E**, Ibal1-positive microglial cells (magenta, indicated by triangles) in the amygdala of control (**D**) and KO (**E**) marmosets. Right, magnified view of rectangle on left. Scale bar, 100 μm (left); 50 μm (right). **F**, Density of microglial cells in the amygdala of marmosets. n = 6 slices, two-tailed unpaired *t*-test, t(10) = 0.7853, *P* = 0.2799. n.s., not significant; data are mean ± SEM. Source data were provided in figure 1-source data 2.

**Figure 2—figure supplement 1.**
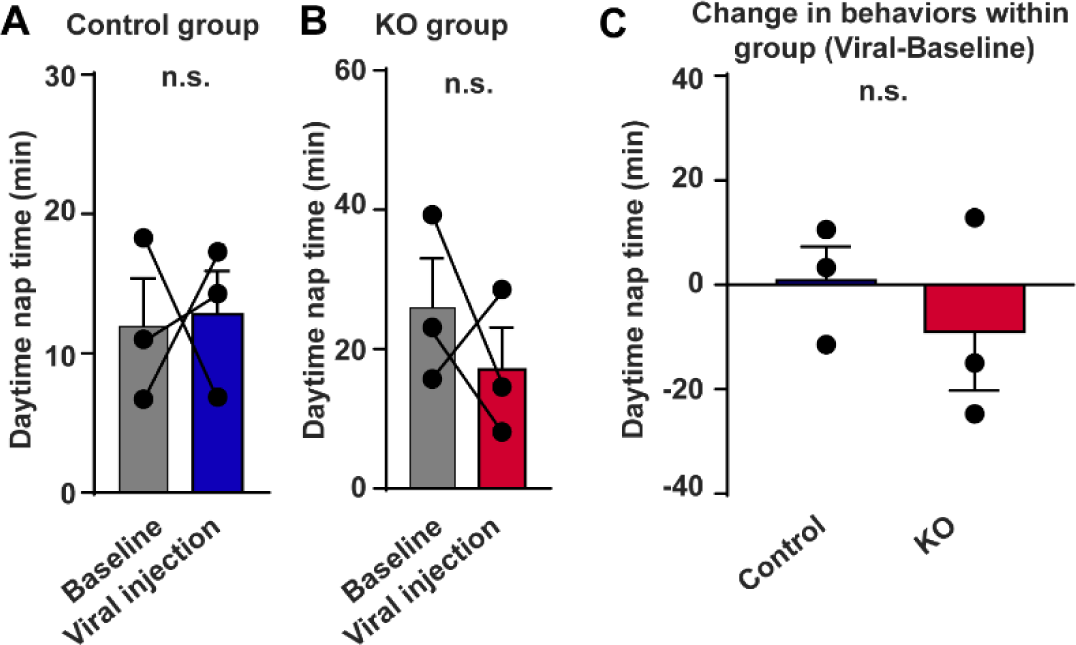
Unaffected daytime nap time after CB1R knockout in the amygdala of adult marmosets. **A**, **B**, Five-day average daytime nap time in control (**A**) and KO (**B**) groups. n = 3, two-tailed paired *t*-test, t(2) = 0.1251, *P* = 0.9119 (**A**); t(2) = 0.7951, *P* = 0.5099 (**B**). **C**, Comparison of changes in daytime nap time between control and KO groups. n = 3, two-tailed unpaired *t*-test, t(4) = 0.7517, *P* = 0.4940. n.s., not significant; data are mean ± SEM. Source data were provided in figure 2-source data 2.

**Figure 2—figure supplement 2.**
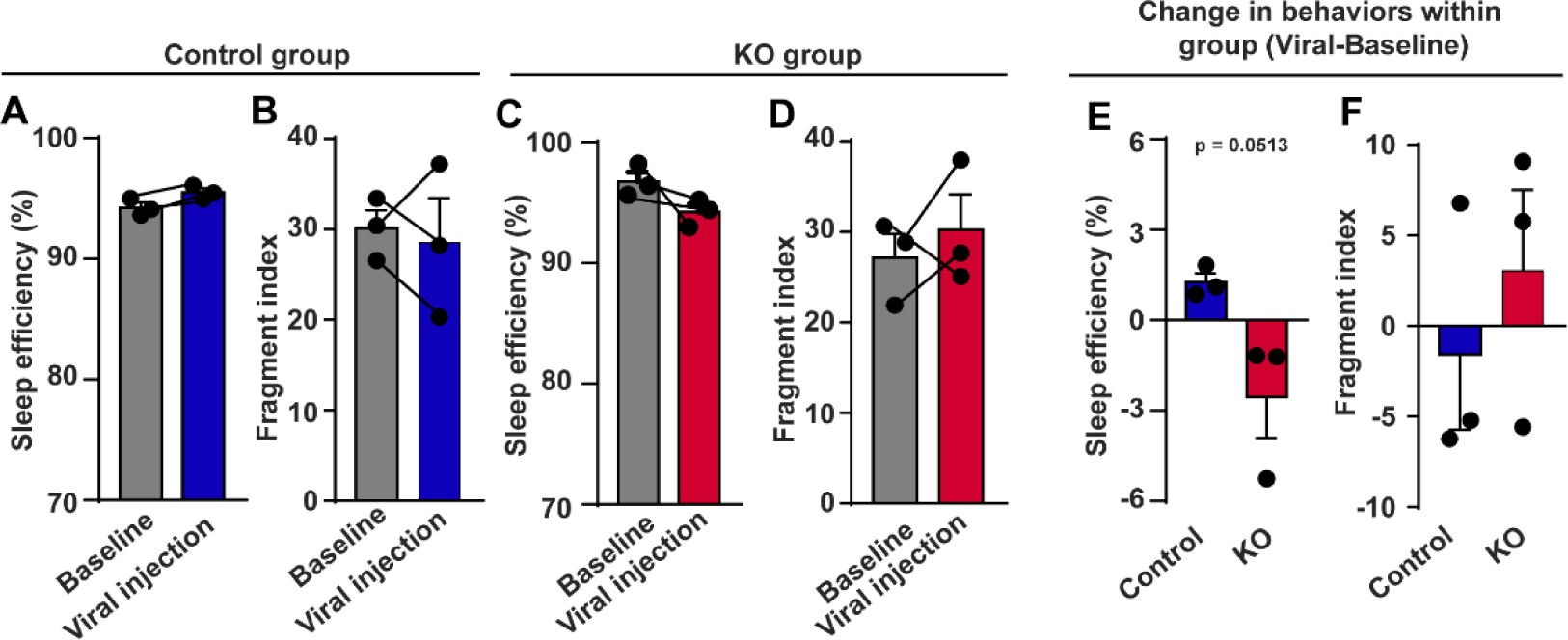
Decreased sleep quality after CB1R knockout in the amygdala of adult marmosets. **A**, **C**, Seven-day average sleep efficiency in control (**A**) and KO (**C**) groups. n = 3, two-tailed paired *t*-test, t(2) = 4.255, *P* = 0.0510 (**A**); t(2) = 1.900, *P* = 0.1978 (**C**). **B**, **D**, Seven-day average sleep fragment index in control (**B**) and KO (**D**) groups. n = 3, two-tailed paired *t*-test, t(2) = 0.3714, *P* = 0.7460 (**B**); t(2) = 0.6975, *P* = 0.5577 (**D**). **E**, **F**, Comparison of changes in sleep efficiency (**E**) and fragment index (**F**) between control and KO groups. n = 3, two-tailed unpaired *t*-test, t(4) = 2.751, *P* = 0.0513 (**E**); t(4) = 0.7624, *P* = 0.4883 (**F**). Data are mean ± SEM. Source data were provided in figure 2-source data 3.

**Figure 3—figure supplement 1.**
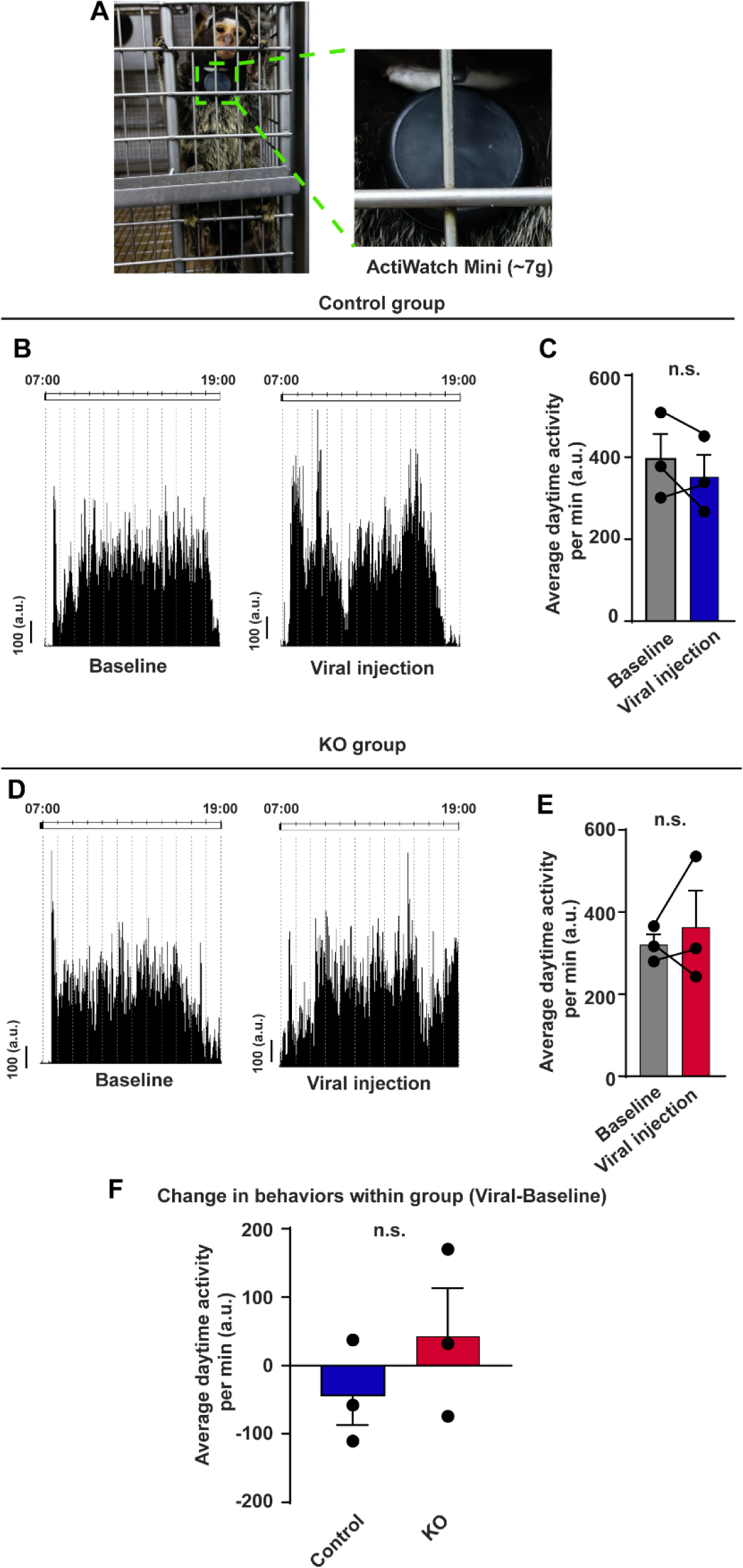
Intact activity in home cage after CB1R knockout in the amygdala of adult marmosets. **A**, Left, representative marmoset wearing ActiWatch Mini. Right, magnified view of ActiWatch Mini in left rectangle. **B**, **D**, Activity counts in home cage before virus injection (left) and after virus injection (right) in a control (**B**) and a KO (**D**) marmoset. **C**, **E**, Seven-day average diurnal activity counts per min in home cage in control (**C**) and KO (**E**) groups. n = 3, two-tailed paired *t*-test, t(2) = 1.006, *P* = 0.4203 (**C**); t(2) = 0.6041, *P* = 0.6072 (**E**). **F**, Comparison of changes in diurnal activity counts per min in home cage between control and KO groups. n = 3, two-tailed unpaired *t*-test, t(4) = 1.041, *P* = 0.3568. n.s., not significant; data are mean ± SEM. Source data were provided in figure 3-source data 2.

**Figure 4—figure supplement 1.**
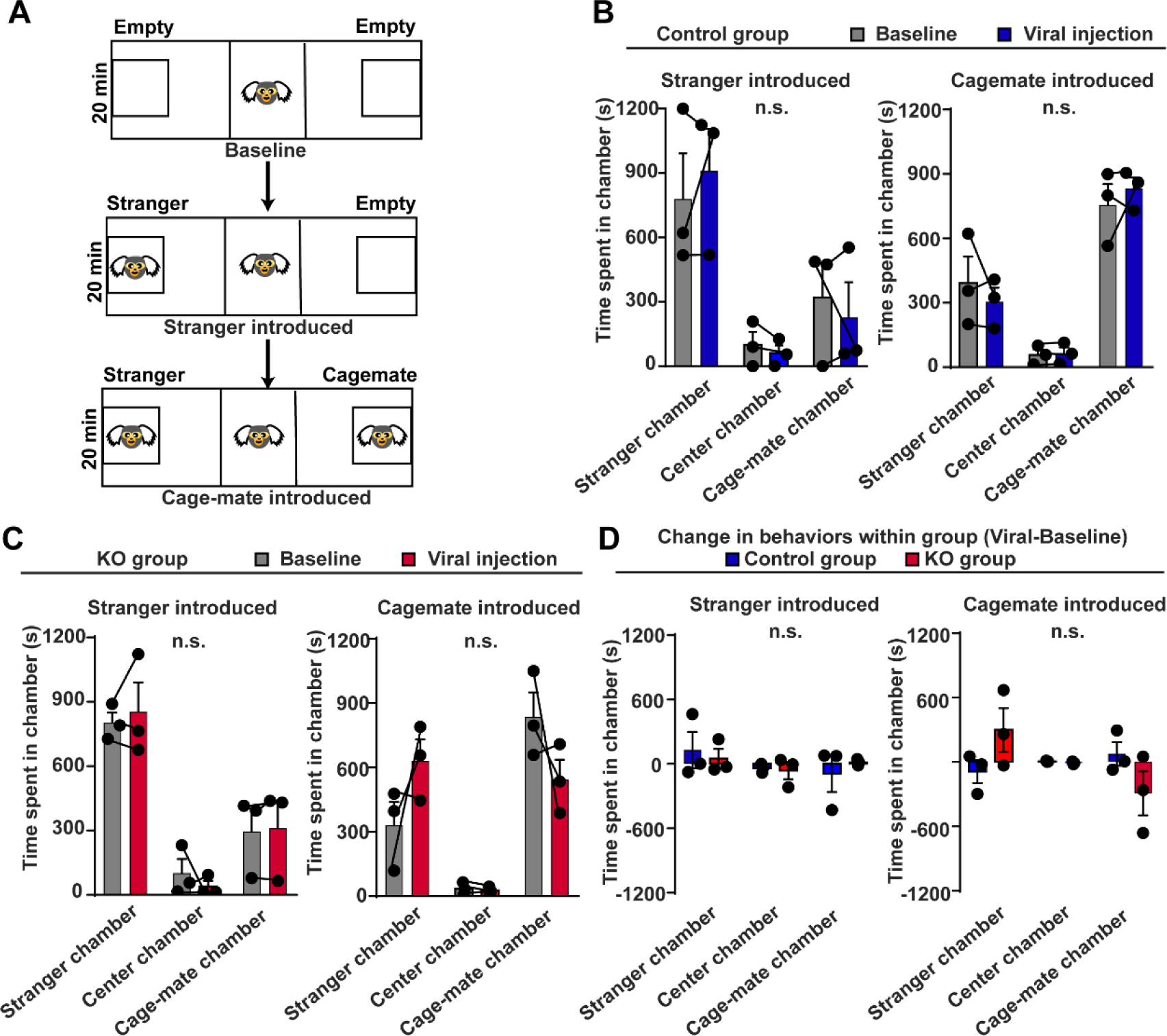
Unaffected social behavior in three-chamber social test after CB1R knockout in the amygdala of adult marmosets. **A**, Paradigm of three-chamber social test in marmosets. **B**, Three-day average of time spent in different chambers during stranger introduced period (left) and cage-mate introduced period (right) in control group. n = 3, Two-way ANOVA, F(2,6) = 0.7025, *P* = 0.5320 (left); F(2,6) = 0.8705, *P* = 0.4657 (right). **C**, Three-day average of time spent in different chambers during stranger introduced period (left) and cage-mate introduced period (right) in KO group. N = 3, Two-way ANOVA, F(2,6) = 0.6798, *P* = 0.5419 (left); F(2,6) = 3.123, *P* = 0.1176 (right). **D**, Comparison of changes in social behavior during stranger introduced period (left) and cage-mate introduced period (right) between control and KO groups. n = 3, Two-way ANOVA, F(2,6) = 0.2880, *P* = 0.7596 (left); F(2,6) = 2.211, *P* = 0.1908 (right). n.s., not significant; data are mean ± SEM. Source data were provided in figure 4-source data 2

**Figure 5—figure supplement 1.**
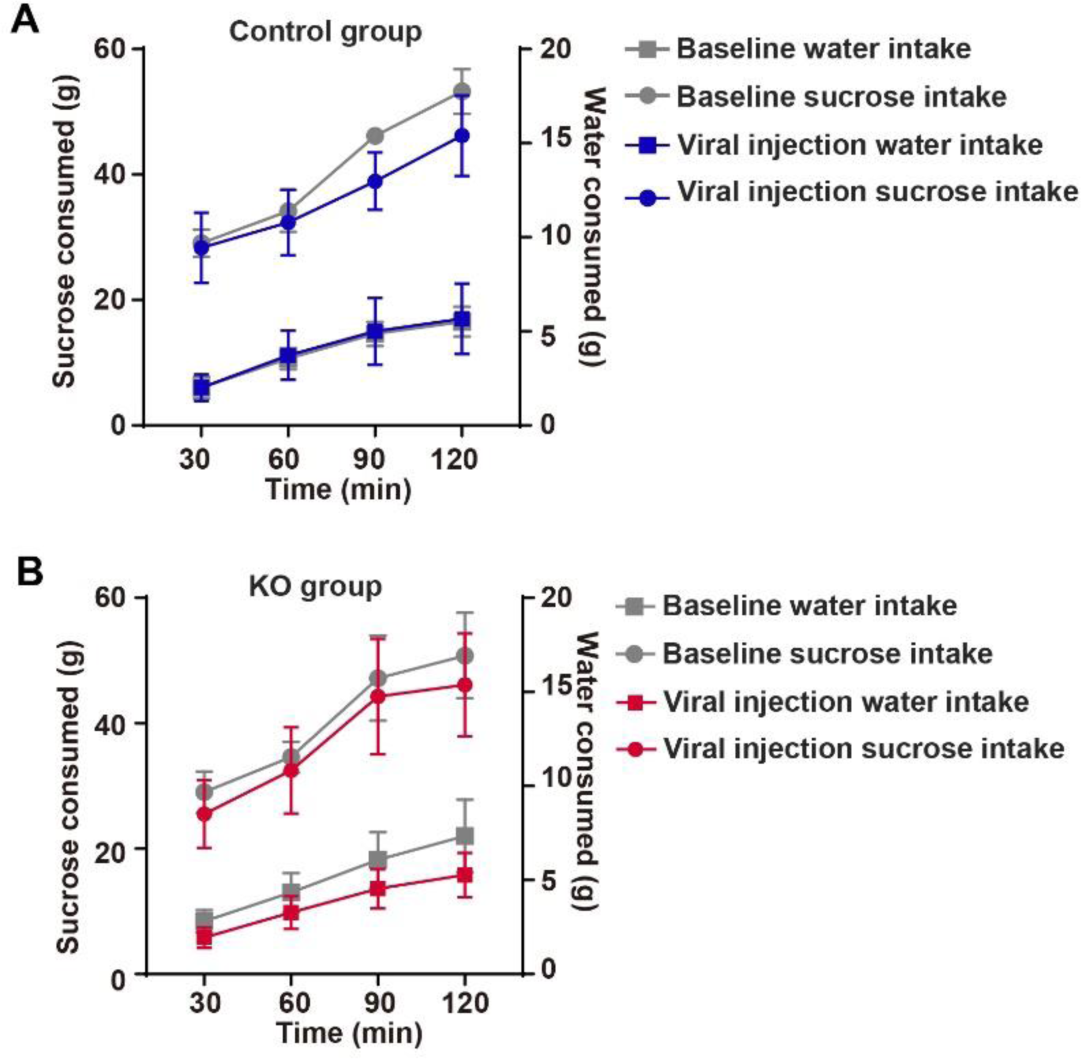
Sucrose preference test. **A**, **B**, Consumption of water and sucrose across 2-h sucrose preference test in control (**A**) and KO groups (**B**). Sucrose consumption is shown on left y-axis and water consumption is shown on right y-axis.

## References

1. Akirav, I. (2013). Cannabinoids and glucocorticoids modulate emotional memory after stress. Neurosci Biobehav Rev 37, 2554–2563.

2. Alexander, L., Gaskin, P.L.R., Sawiak, S.J., Fryer, T.D., Hong, Y.T., Cockcroft, G.J., Clarke, H.F., and Roberts, A.C. (2019). Fractionating Blunted Reward Processing Characteristic of Anhedonia by Over-Activating Primate Subgenual Anterior Cingulate Cortex. Neuron 101, 307–320 e306.

3. Allison, K.C., Spaeth, A., and Hopkins, C.M. (2016). Sleep and Eating Disorders. Curr Psychiatry Rep 18, 92.

4. Bezerra, B.M., and Souto, A. (2008). Structure and usage of the vocal repertoire of Callithrix jacchus. International Journal of Primatology 29, 671.

5. Braesicke, K., Parkinson, J.A., Reekie, Y., Man, M.S., Hopewell, L., Pears, A., Crofts, H., Schnell, C.R., and Roberts, A.C. (2005). Autonomic arousal in an appetitive context in primates: a behavioural and neural analysis. Eur J Neurosci 21, 1733–1740.

6. Calhoon, G.G., and Tye, K.M. (2015). Resolving the neural circuits of anxiety. Nature neuroscience 18, 1394–1404.

7. Camus, S., Ko, W.K.D., Pioli, E., and Bezard, E. (2015). Why bother using non-human primate models of cognitive disorders in translational research? Neurobiology of Learning and Memory 124, 123–129.

8. Castillo, P.E., Younts, T.J., Chavez, A.E., and Hashimotodani, Y. (2012). Endocannabinoid signaling and synaptic function. Neuron 76, 70–81.

9. Chida, Y., and Steptoe, A. (2009). Cortisol awakening response and psychosocial factors: a systematic review and meta-analysis. Biological psychology 80, 265–278.

10. Choi, K., Le, T., McGuire, J., Xing, G., Zhang, L., Li, H., Parker, C.C., Johnson, L.R., and Ursano, R.J. (2012). Expression pattern of the cannabinoid receptor genes in the frontal cortex of mood disorder patients and mice selectively bred for high and low fear. J Psychiatr Res 46, 882–889.

11. Dal Monte, O., Costa, V.D., Noble, P.L., Murray, E.A., and Averbeck, B.B. (2015). Amygdala lesions in rhesus macaques decrease attention to threat. Nature communications 6, 10161.

12. Dunsmoor, J.E., and Paz, R. (2015). Fear Generalization and Anxiety: Behavioral and Neural Mechanisms. Biol Psychiatry 78, 336–343.

13. Feng, G., Jensen, F.E., Greely, H.T., Okano, H., Treue, S., Roberts, A.C., Fox, J.G., Caddick, S., Poo, M.M., Newsome, W.T., et al. (2020). Opportunities and limitations of genetically modified nonhuman primate models for neuroscience research. Proceedings of the National Academy of Sciences of the United States of America 117, 24022–24031.

14. Gothard, K.M. (2020). Multidimensional processing in the amygdala. Nature reviews Neuroscience 21, 565–575.

15. Hamilton, J.P., and Gotlib, I.H. (2008). Neural substrates of increased memory sensitivity for negative stimuli in major depression. Biol Psychiatry 63, 1155–1162.

16. Hanna, R.E., and Doench, J.G. (2020). Design and analysis of CRISPR–Cas experiments. Nature biotechnology 38, 813–823.

17. Haring, M., Kaiser, N., Monory, K., and Lutz, B. (2011). Circuit specific functions of cannabinoid CB1 receptor in the balance of investigatory drive and exploration. PloS one 6, e26617.

18. Hungund, B.L., Vinod, K.Y., Kassir, S.A., Basavarajappa, B.S., Yalamanchili, R., Cooper, T.B., Mann, J.J., and Arango, V. (2004). Upregulation of CB1 receptors and agonist-stimulated [35S]GTPgammaS binding in the prefrontal cortex of depressed suicide victims. Mol Psychiatry 9, 184–190.

19. Hyde, L.W., Gorka, A., Manuck, S.B., and Hariri, A.R. (2011). Perceived social support moderates the link between threat-related amygdala reactivity and trait anxiety. Neuropsychologia 49, 651–656.

20. Izpisua Belmonte, J.C., Callaway, E.M., Caddick, S.J., Churchland, P., Feng, G., Homanics, G.E., Lee, K.F., Leopold, D.A., Miller, C.T., Mitchell, J.F., et al. (2015). Brains, genes, and primates. Neuron 86, 617–631.

21. Janak, P.H., and Tye, K.M. (2015). From circuits to behaviour in the amygdala. Nature 517, 284–292.

22. Jayakar, R., Tone, E.B., Crosson, B., Turner, J.A., Anderson, P.L., Phan, K.L., and Klumpp, H. (2020). Amygdala volume and social anxiety symptom severity: Does segmentation technique matter? Psychiatry Res Neuroimaging 295, 111006.

23. Jennings, C.G., Landman, R., Zhou, Y., Sharma, J., Hyman, J., Movshon, J.A., Qiu, Z., Roberts, A.C., Roe, A.W., Wang, X., et al. (2016). Opportunities and challenges in modeling human brain disorders in transgenic primates. Nature neuroscience 19, 1123–1130.

24. Li, H., Wu, S., Ma, X., Li, X., Cheng, T., Chen, Z., Wu, J., Lv, L., Li, L., Xu, L., et al. (2021). Co-editing PINK1 and DJ-1 Genes Via Adeno-Associated Virus-Delivered CRISPR/Cas9 System in Adult Monkey Brain Elicits Classical Parkinsonian Phenotype. Neurosci Bull 37, 1271–1288.

25. Lutz, B., Marsicano, G., Maldonado, R., and Hillard, C.J. (2015). The endocannabinoid system in guarding against fear, anxiety and stress. Nature reviews Neuroscience 16, 705–718.

26. Marx, V. (2016). Neurobiology: learning from marmosets. Nat Methods 13, 911–916.

27. Melamed, J.L., de Jesus, F.M., Maior, R.S., and Barros, M. (2017). Scopolamine Induces Deficits in Spontaneous Object-Location Recognition and Fear-Learning in Marmoset Monkeys. Front Pharmacol 8, 395.

28. Miller, C.T., Freiwald, W.A., Leopold, D.A., Mitchell, J.F., Silva, A.C., and Wang, X. (2016). Marmosets: A Neuroscientific Model of Human Social Behavior. Neuron 90, 219–233.

29. Moreira, F.A., Grieb, M., and Lutz, B. (2009). Central side-effects of therapies based on CB1 cannabinoid receptor agonists and antagonists: focus on anxiety and depression. Best practice & research Clinical endocrinology & metabolism 23, 133–144.

30. Morrison, S.E., and Salzman, C.D. (2010). Re-valuing the amygdala. Current opinion in neurobiology 20, 221–230.

31. Ohno-Shosaku, T., and Kano, M. (2014). Endocannabinoid-mediated retrograde modulation of synaptic transmission. Current opinion in neurobiology 29, 1–8.

32. Okano, H. (2021). Current Status of and Perspectives on the Application of Marmosets in Neurobiology. Annual review of neuroscience 44, 27–48.

33. Okano, H., Sasaki, E., Yamamori, T., Iriki, A., Shimogori, T., Yamaguchi, Y., Kasai, K., and Miyawaki, A. (2016). Brain/MINDS: A Japanese National Brain Project for Marmoset Neuroscience. Neuron 92, 582–590.

34. Paquet, J., Kawinska, A., and Carrier, J. (2007). Wake detection capacity of actigraphy during sleep. Sleep 30, 1362–1369.

35. Park, S.C., and Kim, Y.K. (2020). Anxiety Disorders in the DSM-5: Changes, Controversies, and Future Directions. Advances in experimental medicine and biology 1191, 187–196.

36. Penninx, B.W.J.H., Pine, D.S., Holmes, E.A., and Reif, A. (2021). Anxiety disorders. The Lancet 397, 914–927.

37. Qiu, P., Jiang, J., Liu, Z., Cai, Y., Huang, T., Wang, Y., Liu, Q., Nie, Y., Liu, F., Cheng, J., et al. (2019). BMAL1 knockout macaque monkeys display reduced sleep and psychiatric disorders. National Science Review 6, 87–100.

38. Ramikie, T.S., and Patel, S. (2012). Endocannabinoid signaling in the amygdala: anatomy, synaptic signaling, behavior, and adaptations to stress. Neuroscience 204, 38–52.

39. Rein, B., Ma, K., and Yan, Z. (2020). A standardized social preference protocol for measuring social deficits in mouse models of autism. Nat Protoc 15, 3464–3477.

40. Rey, A.A., Purrio, M., Viveros, M.P., and Lutz, B. (2012). Biphasic effects of cannabinoids in anxiety responses: CB1 and GABA(B) receptors in the balance of GABAergic and glutamatergic neurotransmission. Neuropsychopharmacology 37, 2624–2634.

41. Roche, M., O’Connor, E., Diskin, C., and Finn, D.P. (2007). The effect of CB(1) receptor antagonism in the right basolateral amygdala on conditioned fear and associated analgesia in rats. Eur J Neurosci 26, 2643–2653.

42. Ross, C.N., Adams, J., Gonzalez, O., Dick, E., Giavedoni, L., Hodara, V.L., Phillips, K., Rigodanzo, A.D., Kasinath, B., and Tardif, S.D. (2019). Cross-sectional comparison of health-span phenotypes in young versus geriatric marmosets. Am J Primatol 81, e22952.

43. Shen, C.J., Zheng, D., Li, K.X., Yang, J.M., Pan, H.Q., Yu, X.D., Fu, J.Y., Zhu, Y., Sun, Q.X., Tang, M.Y., et al. (2019). Cannabinoid CB(1) receptors in the amygdalar cholecystokinin glutamatergic afferents to nucleus accumbens modulate depressive-like behavior. Nat Med 25, 337–349.

44. Spasojevic, N., Stefanovic, B., Jovanovic, P., and Dronjak, S. (2016). Anxiety and Hyperlocomotion Induced by Chronic Unpredictable Mild Stress Can Be Moderated with Melatonin Treatment. Folia biologica 62, 250–257.

45. Tovote, P., Fadok, J.P., and Luthi, A. (2015). Neuronal circuits for fear and anxiety. Nature reviews Neuroscience 16, 317–331.

46. Wellman, L.L., Forcelli, P.A., Aguilar, B.L., and Malkova, L. (2016). Bidirectional Control of Social Behavior by Activity within Basolateral and Central Amygdala of Primates. The Journal of neuroscience : the official journal of the Society for Neuroscience 36, 8746–8756.

47. Wu, S.H., Li, X., Qin, D.D., Zhang, L.H., Cheng, T.L., Chen, Z.F., Nie, B.B., Ren, X.F., Wu, J., Wang, W.C., et al. (2021). Induction of core symptoms of autism spectrum disorder by in vivo CRISPR/Cas9-based gene editing in the brain of adolescent rhesus monkeys. Science Bulletin 66, 937–946.

48. Zhou, Y., Sharma, J., Ke, Q., Landman, R., Yuan, J., Chen, H., Hayden, D.S., Fisher, J.W., 3rd, Jiang, M., Menegas, W., et al. (2019). Atypical behaviour and connectivity in SHANK3-mutant macaques. Nature 570, 326-331.

